# Dynamic signatures of the Eureka effect: An EEG study

**DOI:** 10.1101/2021.02.27.433162

**Authors:** Yiqing Lu, Wolf Singer

**Affiliations:** Ernst Strüngmann Institute for Neuroscience in Cooperation with Max Planck Society, 60528 Frankfurt am Main, Germany; Frankfurt Institute for Advanced Studies, 60438 Frankfurt am Main, Germany; Department of Neurophysiology, Max Planck Institute for Brain Research, 60528 Frankfurt am Main, Germany; Department of Biology, Technische Universität Darmstadt, 64287 Darmstadt, Germany

**Keywords:** Eureka effect, EEG, coherence, phase locking, lateralization, dimensionality

## Abstract

The Eureka effect refers to the common experience of suddenly solving a problem. Here we study this effect in a pattern recognition paradigm that requires the segmentation of complex scenes and recognition of objects on the basis of Gestalt rules and prior knowledge. In the experiments both sensory evidence and prior knowledge were manipulated in order to obtain trials that do or do not converge towards a perceptual solution. Subjects had to detect objects in blurred scenes and signal recognition with manual responses. Neural dynamics were analyzed with high-density Electroencephalography (EEG) recordings. The results show significant changes of neural dynamics with respect to spectral distribution, coherence, phase locking, and fractal dimensionality. The Eureka effect was associated with increased coherence of oscillations in the alpha and theta band over widely distributed regions of the cortical mantle predominantly in the right hemisphere. This increase in coherence was associated with a decrease of beta band activity over parietal and central regions, and with a decrease of alpha power over frontal and occipital areas. In addition, there was a lateralized reduction of fractal dimensionality for activity recorded from the right hemisphere. These results suggest that the transition towards the solution of a perceptual task is mainly associated with a change of network dynamics in the right hemisphere that is characterized by enhanced coherence and reduced complexity. We propose that the Eureka effect requires cooperation of cortical regions involved in working memory, creative thinking, and the control of attention.

## Introduction

Nervous systems need to be able to distinguish activity patterns associated with the search for the solution of a computational problem from those representing a result. This distinction is necessary in order to terminate further search, to eventually convert the result into action, to permit synaptic plasticity for the storage of results, to allow initiation of a new search process, and to permit access of results to awareness. Humans experience solution states as rewarding and are capable of judging the reliability of a particular result. It is pleasant to suddenly arrive at the solution of a perceptual problem, to understand a previously incomprehensible concept, or to solve a puzzle. This sudden transition from searching for a solution to having found the solution has been termed the Eureka effect (Kaplan and Simon, 1990; Sternberg and Davidson, 1995; Ahissar and Hochstein, 1997; Kounios and Beeman, 2014; Sprugnoli et al., 2017). Some of the features of the Eureka experience are probably also shared by higher mammals (Tovee et al., 1996).

However, very little is known about the neuronal underpinnings of the Eureka effect. One of the reasons is that it is hard to predict under which circumstances and exactly when the Eureka effect occurs. Investigations of the Eureka effect are fraught with several methodological problems. First, measurements need to be performed with high temporal resolution because the moment when the Eureka effect occurs is often unpredictable. Second, neurophysiological studies require a sufficient number of comparable trials for analysis.

In the present study, we applied black-and-white Mooney images (Dolan et al., 1997; Tallon-Baudry and Bertrand, 1999; McKeeff and Tong, 2007; Giovannelli et al., 2010; Castelhano et al., 2013) as the paradigm to induce the Eureka effect. The most interesting feature of Mooney images is that they are initially difficult to interpret, but can eventually lead to a stable percept of objects. This percept can also be facilitated by cues, which usually consist of original images (color or greyscale images) of the Mooney images (Hsieh et al., 2010; Goold and Meng, 2016). Mooney images have several advantages as test material. First, they permit precise manipulation of parameters such as luminance, contrast, size, and pixels. Second, Mooney images allow the combination of natural objects with complex backgrounds. This permits the induction of strong Eureka effects and the search for neuronal correlates. To maximize the occurrence of Eureka experiences we developed a method to generate and fine-tune Mooney images that evoke Eureka experiences reliably and reproducibly.

How are ambiguous inputs from Mooney images transformed to unambiguous perceptual solutions? It is suggested that prior experience, stored in memory, is used to modulate peripheral processing in order to facilitate scene segmentation, perceptual grouping, and recognition (Dolan et al., 1997; Gorlin et al., 2012). The Bayesian hypothesis of perception posits that internal information, made available by top-down processes is matched with sensory evidence (Kersten et al., 2004; Friston and Stephan, 2007; Clark, 2013; de Lange et al., 2018). Thus, the brain is assumed to perform a probabilistic inference that can optimize sensory information to minimize the mismatch between internal and external information. This function can be interpreted as perceptual closure which consists of the completion of incomplete sensory evidence and the correct binding of components into a coherent percept (Grutzner et al., 2010; Moratti et al., 2014). One proposal is that this is achieved through synchronization to transiently form a coherently active assembly of neurons that generates the perception of a complete object (Singer and Gray, 1995).

Assuming that synchronized activities within and across cortical areas are associated with feature integration and perception and that synchronization is facilitated by engagement of neuronal populations in oscillatory activity (Gray et al., 1989; Singer, 1999; Sehatpour et al., 2008; Fries, 2009; Hipp et al., 2011; Volberg et al., 2013; Singer, 2018), we wondered whether and if so, how oscillatory and synchronized activities are involved in the Eureka effect. We hypothesized that solution states might be associated with enhanced coherence of neuronal activity as they are likely to result from successful integration of distributed computational results. To examine this hypothesis, we presented Mooney stimuli to induce the Eureka effect, captured neural activity with high-density EEG recordings and then investigated how these neuronal responses were modulated during the Eureka effect. As the brain can be considered as a complex system with non-linear dynamics, parameters taken from complexity theory should also be indicators of state transitions (Sporns et al., 2004; Bullmore and Sporns, 2012; Avena-Koenigsberger et al., 2017). For this reason, we measured not only the power and coherence of oscillatory activity but also the fractal dimensionality of activity vectors (Nikolic et al., 2008; Singer and Lazar, 2016) to further characterize the phase transitions associated with the Eureka effect.

## Materials and Methods

### The selection of images

The original images used for the construction of the stimuli were taken from the Caltech-256 Object Category Dataset (Griffin et al., 2007) and Flickr. More than 30,000 images were included in our initial database. We then manually chose images for further testing according to the following criteria. 1) Familiarity: we chose well known objects, such as animals, plants, furniture, tools, instruments, vehicles, and so on, 2) Natural and complex background: the images were taken from natural scenes; the target objects being embedded in a complex background, which can render identification difficult.

### The procedure of manipulating the images

The original images were first converted to 256-level greyscale images. As the distribution of greyscale pixels varied widely and was non-normal and as luminance levels and contrast also differed, adjustments were required. Luminance and contrast were equated with histogram equalization which generates even contrast distributions (Lim, 1990).

To create Mooney images with different degradation levels, we applied a frequency-domain Gaussian filter on the greyscale images processed according to the methods described above (Haddad and Akansu, 1991). Lowering the cut-off frequency of the filter increases the difficulty to recognize the object in an image. The filters were generated in three processing steps: 1) The spatial domain of the original image *f(x,y)* is transformed to the frequency domain *F(u,v)* by Fourier transformation; 2) this *F(u,v)* is low-pass filtered with a set of cut-off frequencies to obtain a band passed representation *G(u,v);* 3) this representation *G(u,v)* is then transformed back to the spatial domain *g(x,y)* (i.e. the blurred image) by an inverse Fourier transformation. In order to obtain images with different degradation levels, the low-pass filter was varied between cut-off frequencies ranging from 10 Hz to 80 Hz (10 Hz per interval) (Fig. S1). The median grey level of each image was then chosen as the divide for the assignment of black or white pixels in order to obtain two-tone Mooney images. Subsequently the recognizability of Mooney images with different cut-off frequencies was tested. This led to the selection of the 20 Hz cut-off frequency for the induction of the Eureka effect because it assured transitions from search to solution most reliably (details in Supplementary material).

### Participants

In the pilot experiment aimed at the determination of the cut-off frequency, 10 subjects were included. None of these subjects participated in the main experiment (The detailed information of the pilot experiment is in Supplementary material). Another 29 healthy subjects (Age 25.2 ± 4.0 y, 16 males, 13 females) took part in the main experiment that comprised both behavioral assessment and EEG recordings. None of these subjects participated in the pilot experiment. Twenty-five subjects were included and 4 were rejected due to an insufficient number of correct responses. In addition, EEG data from three of these 25 subjects had to be excluded from EEG analysis due to insufficient valid data because of artifact rejection (The artifact rejection is described in the “EEG data pre-processing” section). All subjects were naïve to the experiment, were righthanded, had normal or corrected-to-normal vision, and had no history of neurological or psychiatric disorders. They gave written informed consent before the experiment. The study was approved by the ethical committee of the Goethe University, Frankfurt, and was conducted in accordance with the Declaration of Helsinki. The subjects were recruited from local universities and got paid 15 Euros per hour for their participation.

### Visual stimuli

Presentation^®^ (V10.3, Neurobehavioral Systems) was used for stimulus presentation and response collection. All stimuli were generated using Matlab (The Mathworks). The stimuli were displayed as 150 x 150 pixels matrices at the center of a monitor screen with a refresh rate of 60 Hz and located 70 cm from the subjects’ eye plane, subtending a visual angle of 4.4° x 4.4°, surrounded by grey background, with a grey level of 0.5 on a greyscale of [0, 1].

### The task during EEG recording

For each subject, 160 different stimuli were used. At the beginning of each trial, a fixation cross was presented on grey background with a randomized duration between 2 s and 3 s. Then, a Mooney image was displayed for 8 s. We address this presentation as the *first stage* of a trial. If the subject could identify the object in the image, they had to press the button “Yes” as soon as possible during this 8 s interval which terminated the trial. If they could not identify the object in the 8 s, the Mooney image disappeared. Then, a greyscale image was provided as the cue after a random interval of 1.5 - 2 s. This image lasted for 4 s. In 50% of the trials, the greyscale images were congruent with the last Mooney images and served as cues for the subsequent identification. In the other 50% of the trials, they were incongruent. We address this greyscale image presentation as the *second stage* of a trial. Once the greyscale image had disappeared, the last Mooney image appeared again after a random interval of 2 - 3 s. We address this repeated Mooney image presentation as the *third stage* of a trial. At this stage, subjects had to press the button “Yes” or “No”, indicating whether or not they could identify the object in the Mooney image. If subjects responded with a prompt “Yes” in the matching trials, we took this as evidence that they had experienced a Eureka effect, the sudden recognition of a pattern that they had been unable to identify in the *first stage.* Figure 1 illustrates the task procedure. The order of trials and the order of experimental conditions were randomized. The experiment was evenly divided into four blocks. A break of 2 - 3 minutes was introduced after each block. Prior to the experiment, a training session with a different set of images was performed in order to allow each subject to practice.

**Figure 1.**
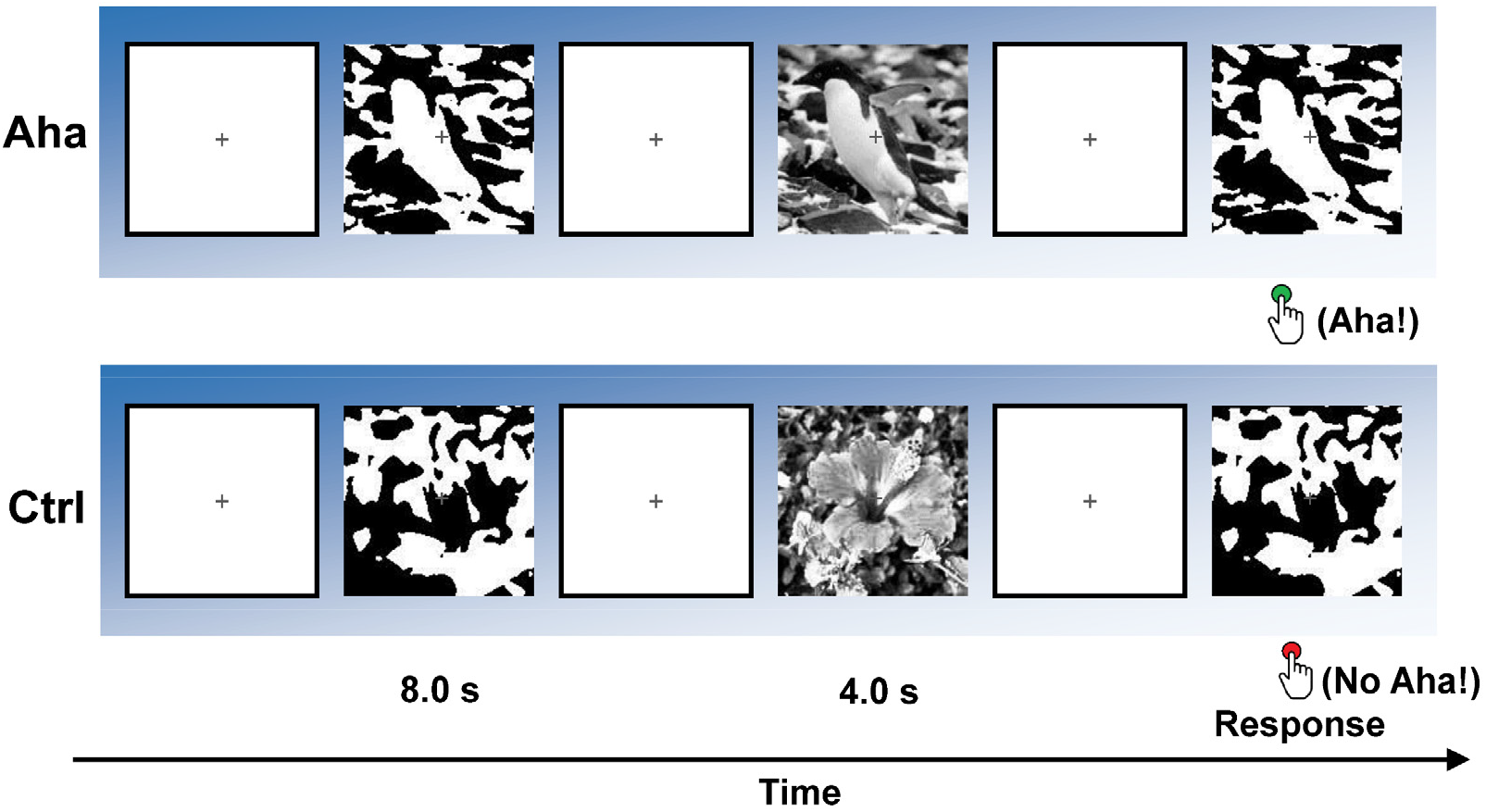
The paradigm during EEG recording. The upper row shows an example of Eureka (Aha) trials which were composed of Mooney images and their congruent greyscale images. The lower row shows an example of Control (Ctrl) trials which are composed of Mooney images and incongruent greyscale images.

### Data acquisition

The experiment was implemented in an electrically shielded, sound-attenuated, darkroom. Subjects watched the monitor which was outside the room through an electrically shielded window. The EEG was recorded with a HydroCel Geodesic Sensor Net 130 (HCGSN) with 128 channels (Electrical Geodesics, Inc.), with the reference electrode at Cz. The electrodes were spaced over the head following the instructions. Electrode impedances were kept < 50 kΩ. Data were sampled at 1000 Hz and digitally saved on a Mac system for off-line analysis. The data used in this study are available via a request to the corresponding author upon signing a formal data sharing agreement.

### Behavioral analysis

We subdivided trials into two groups: an Eureka (Aha) group and a control (Ctrl) group. The Aha group consisted of trials (with congruent cue) and “Yes” responses at the *third stage.* The Ctrl group consisted of trials (with incongruent cue) and “No” responses at the *third stage.* Trails that did not meet these requirements (9.5% of trials) were discarded from further analysis. We then compared the distributions of reaction times (RTs) for trials of Aha and Ctrl groups.

### EEG data pre-processing

In Net Station software 4.5.1 (Electrical Geodesics, Inc.), the continuous EEG signal was high-pass filtered (0.3 Hz) and notch filtered (50 Hz), and then segmented into a series of 2 – 10 s long epochs (depend on the reaction time). Each trial segment started 1 s before the onset of the *third stage* and ended 1 s after the response during the *third stage*. In the following steps, data were analyzed using the Matlab toolbox FieldTrip (Oostenveld et al., 2011) and our customized scripts. The semi-automatic artifact rejection was supplemented by visual inspection, Fieldtrip scripts, and independent component analysis (ICA) to detect electrode drifts, eye movements, electromyographic and electrocardiographic interference (Bell and Sejnowski, 1995; Amari et al., 1996; Lee et al., 1999; Anemuller et al., 2003). Rejected channels were interpolated using spherical splines. To avoid contamination of neural activity induced by stimulus onset rather than Eureka effects, only the trials with reaction times longer than 0.5 s were analyzed.

### Spectral analysis

To investigate the time-frequency activities that are involved in the Eureka effect, we analyzed the spectral changes of EEG signals over the whole electrode space, using the Hanning taper approach, with a 5-cycle time window and variable width of frequency smoothing depending on the frequency, with 1 Hz steps. The analysis time window was 1 s (for stimulus-locked epochs, from −0.25 to 0.75 s; for response-locked epochs, from −0.75 to 0.25 s). For baseline correction, a time period after stimulus onset of the *first stage* was used that had the same length as the analysis window used in the *third stage,* whose duration was determined by the respective reaction time. In summary, the baseline correction and the comparison between Aha and Ctrl can be described as a simple formula: *Diff = (M3_Aha_ – M1_Aha_) - (M3_ctrl_ – M1_ctrl_)*, in which, *Diff* is the difference between Aha and Ctrl; *M3* and *M1* refer to data from Mooney images at the *third stage* and the *first stage* respectively. The subtraction removes the components resulting from the stimulus-evoked responses. Finally, we calculated grand averages for each condition and participant. The theta, alpha, beta, and gamma bands were defined by the following frequency ranges, respectively, 4–7 Hz, 8–12 Hz, 13–30 Hz, and 31–100 Hz.

### Coherence

Brain networks are defined both anatomically and functionally. Given the high degree of anatomical connectedness among processing streams, any cognitive and executive task requires fast and flexible formation of functional networks. One suitable measure for functional connectivity is coherence (Rosenberg et al., 1989). It takes into account both phase and amplitude components of signals and provides information about the functional coupling of network nodes. We therefore calculated coherence in specific frequency bands in selected time windows between electrode positions/channels.

In order to avoid the bias that may be introduced by unequal numbers of trials, the number of trials for Aha and Ctrl groups was equalized before comparison by randomly discarding trials from the condition with a larger number of trials. Coherence was calculated according to the formula below:

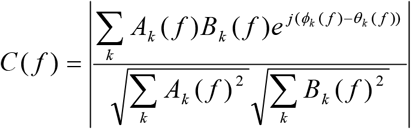

where *A_k_* (*f*)*e*^*jϕ_k_*(*f*)^ and *B_k_* (*f*)*e*^*jθ_k_*(*f*)^ describe the Fourier transformed signals, and *k* the trial number (Srinath and Ray, 2014). The coherence values are real-valued numbers from 0 to 1. One indicates that two signals have perfect coupling, while zero indicates independence. The coherence values described in the results were all baseline corrected according to the method applied for the spectral analysis.

### Phase locking value (PLV)

As coherence values are sensitive to amplitude variations, we also calculated phase locking values. These values reflect phase correlations between two oscillatory signals (Lachaux et al., 1999; Srinath and Ray, 2014) and serve as an index for the synchronization of different neuron groups. The key factor distinguishing the PLV from coherence is that the PLV does not take amplitude into account. It only measures the phase component. Similar to coherence, the PLV also has a real-valued number ranging from 0 to 1. The value 1 indicates that two signals are strictly phase locked, while 0 indicates that their phase relations are random. The PLV can be represented as

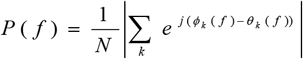

where *e*^*jϕ_k_*(*f*)^ and *e*^*jθ_k_*(*f*)^ are the imaginary parts of the signals, *N* is the number of trials, and *k* is the trial number. The PLV values described in the results are all baseline corrected according to the method applied for spectral analysis.

### Laterality index (LI)

To capture lateralization effects, we calculated laterality indices (LI) (Jansen et al., 2006; Wilke and Schmithorst, 2006; Seghier, 2008). LIs were calculated by evaluating differences between the right and left hemispheres. We used the following formula to determine the LI:

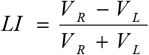

where *V_R_* and *V_L_* refer to averaged values within the right and left hemisphere, or the right and left electrode cluster. A positive or negative value of LI indicates lateralization of neural activity to the right or left side, respectively. To calculate the LI of coherence and PLV, electrodes (Fp1, Fp2, F7, F3, Fz, F4, F8, T3, C3, CPz, C4, T4, T5, P3, Pz, P4, T6, O1, Oz, and O2) were selected according to the 10-20 EEG system.

### Dimensionality

In order to assess changes in the complexity of network dynamics, we determined the fractal dimensionality of the recorded high dimensional time series. The EEG data were normalized to *z*-scores and then analyzed for fractal structure. The principle of the analysis is based on scale-versus-count relationships for dimension *D,* which can be given by

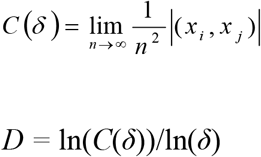

where *n* is the number of measurements, *(x_i_, x_j_)* are pairs of points (|*x_i_* - *x_j_* | < *δ*, *i ≠ J*), *δ* is spheres of a certain size and *C(δ)* is the count of the size (Grassberger and Procaccia, 2004).

The dimensionality was calculated for clusters of electrodes that were selected depending on their positions. Ten clusters were formed according to the proposal by Marie and Trainor (2013). The clusters cover left frontal (LF), left central (LC), left parietal (LP), left temporal (LT), left occipital (LO), right frontal (RF), right central (RC), right parietal (RP), right temporal (RT), and right occipital (RO) areas (Fig. 8, 9). Each cluster included 9 or 10 electrodes.

### Statistics

To evaluate the significance of differences between trials with the Eureka effect and the control condition, we performed a Wilcoxon signed-rank test when samples were not normally distributed; or a *t*-test (two-tailed) when samples were normally distributed. A cluster-based nonparametric randomization test was used to solve the multiple comparisons problem (MCP) (Maris and Oostenveld, 2007). For each paired sample, an independent samples *t*-test was computed. All samples with a *p*-value lower than threshold 0.05 were selected and then clustered on the basis of spatial and temporal adjacency. The sum of the clustered *t* values was calculated. These samples were then randomized across Aha and Ctrl conditions, and the cluster *t* value was analyzed. This step was repeated 1000 times. We then obtained a new distribution of cluster *t* values. On the basis of this distribution, the *t* value from the original data was evaluated. Here, we used the threshold 0.05 for significance and to obtain significant data cluster(s) (for example: including electrodes and time windows for predefined frequencies). Similarly, the cluster-based nonparametric randomization test was also used to correct the *z* value of the Wilcoxon signed-rank test.

## Results

### Behavioral data

Subjects continued 83.9% of the trials till the *third stage* and 90.5% of these trials had correct responses. Correct meant “Yes” response to congruent trials or “No” responses to incongruent trials. This result suggests that the subjects really experienced the Eureka effect and that the comparison between Aha and Ctrl conditions was valid.

The reaction times (RTs) of Eureka (Aha) and Control (Ctrl) trials are summarized in Figure 2. The RTs in Aha trials are significantly faster than the RTs in Ctrl trials (*p* = 3.2229e-05, Wilcoxon signed-rank test). The smoothed distribution of RTs shows that the peak of the Aha RTs distribution is at 0.90 s, while the peak of the Ctrl RTs distribution is at 1.50 s. The Aha RTs distribution is narrower than the Ctrl RTs distribution. The median of the Aha trial RTs is 1.13 s, and that of the Ctrl trials is 2.00 s. In 92.9% of the Aha trials and 99.3% of the Ctrl trials, the RTs were longer than 0.50 s. These trials were kept for further analysis.

**Figure 2.**
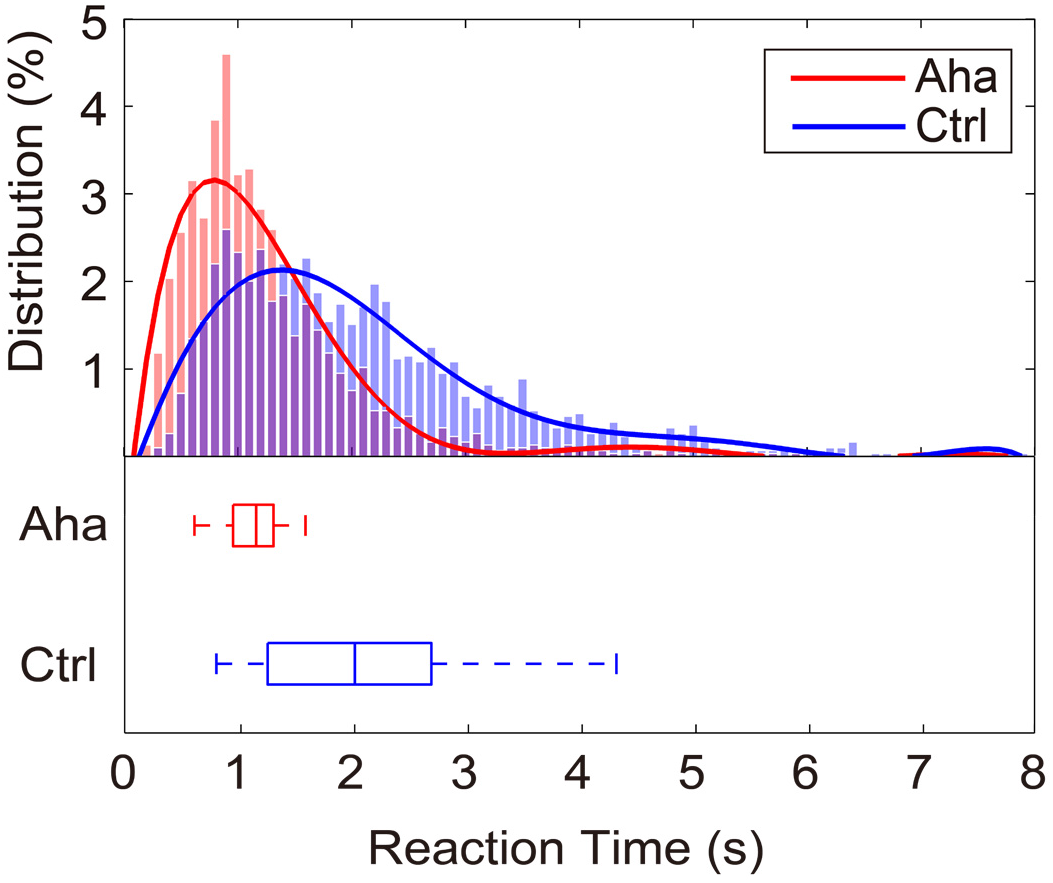
The distribution of reaction times (RTs). Upper panel: distribution of RTs for all trials. Abscissa: RTs, bin width 0.10 s. Ordinate: Frequency of reaction times in %. Red: Aha condition; Blue: Control condition. Continuous lines: Fitted curves. Lower panel: box plots of the respective RTs distributions, with Median values, the 25^th^, and the 75^th^ percentiles. The whiskers indicate extrema.

### The increase of alpha and theta coherence in the right hemisphere

#### Alpha Coherence

To analyze synchronization between channels in the low-frequency range, we calculated coherence in a 500 ms long window starting at stimulus onset (from 0 ms to 500 ms, “stimulus-locked epochs”). This analysis revealed an increase of coherence in the alpha band, which was lateralized and particularly prominent in the right hemisphere, for the Aha condition (Fig. 3A). To quantify this lateralization, we calculated the laterality index (LI) of alpha coherence for channel pairs. This confirmed a significant right hemisphere increase of alpha coherence in the time window from 345 ms to 450 ms post-stimulus *(p* < 0.05), for Aha as compared to Ctrl. Due to the variable reaction time across trials, we also aligned the analysis window to the responses (from −500 ms to response onset, “response-locked epochs”). The right hemisphere lateralized alpha coherence was present also in this response segment and significant in the time window from −355 ms to −275 ms (*p* < 0.05) (Fig. 3B).

**Figure 3.**
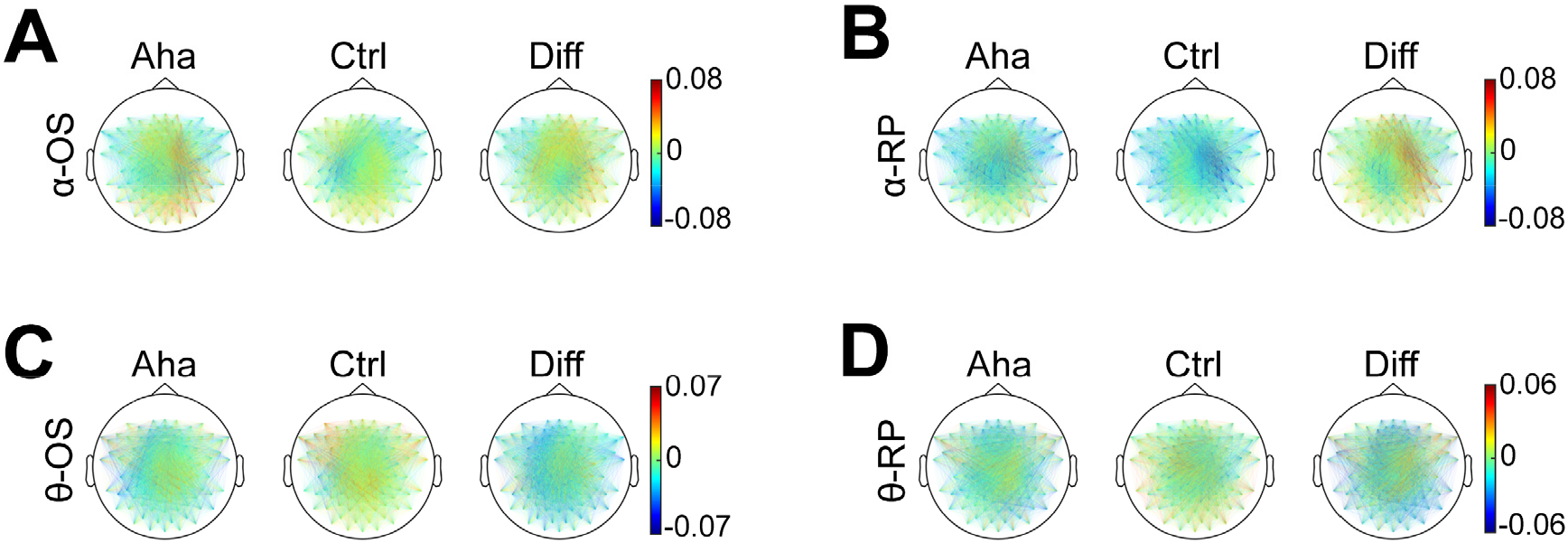
The topographic distribution of alpha (A, B) and theta (C, D) coherence. Baseline corrected. Diff indicates the difference of values between Aha and Ctrl conditions (Aha minus Ctrl). OS represents stimulus onset locked epochs (A, C) (from stimulus onset to 500 ms after onset). RP represents response locked epochs (from 500 ms before response to response). The colored lines represent the coherence values between electrode pairs (coherence values are indicated by the color scales on the right). Note the different (only for clearer illustration) scales in C and D.

#### Theta Coherence

Coherence analysis in the theta band also revealed a right-lateralized increase for the interval 0 - 500 ms for the Aha condition (Fig. 3C). The LI analysis showed that this right lateralization was significant in two time windows: from 125 ms to 260 ms *(p* < 0.05) and from 375 ms to 395 ms *(p* < 0.05), for Aha as compared to Ctrl. For response-locked epochs, there was only a trend for a right hemispheric increase of coherence but this interhemispheric difference was not significant (Fig. 3D).

### The increase of alpha and theta phase locking in the right hemisphere

#### Alpha PLV

To assess changes in phase locking associated with the Eureka effect, we analyzed the phase locking values (PLV) for channel pairs in the alpha band from 0 ms to 500 ms after stimulus onset. Alpha band PLV also increased during the Eureka effect and the topography (Fig. 4A) of this increase resembles that of increases in alpha coherence. The right hemisphere increase of PLVs was significant within the interval from 385 ms to 445 ms post-stimulus *(p* < 0.05), for Aha as compared to Ctrl. This window overlaps with the interval of enhanced coherence (345 - 450 ms). PLVs were also enhanced in the response-locked epochs, again more on the right than the left side but this interhemispheric difference did not reach significance (Fig. 4B).

**Figure 4.**
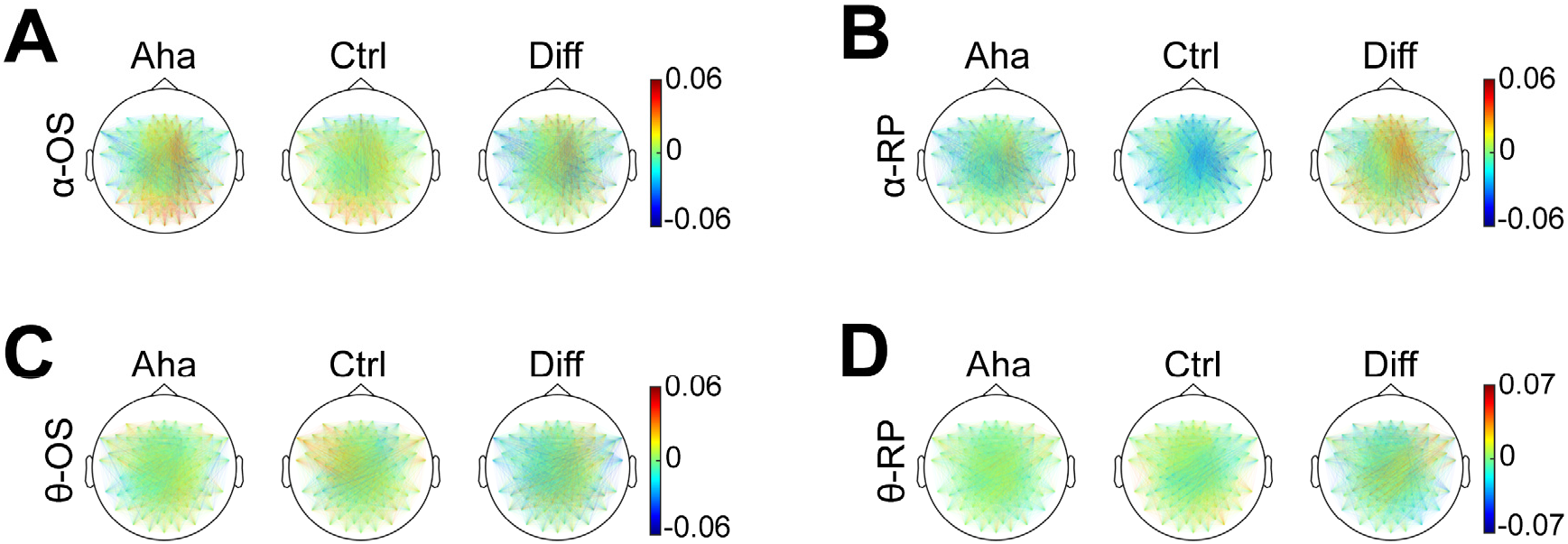
The topographic distribution of alpha and theta PLV. Conventions as in Fig 3. The colored lines represent the PLV values. Note the different (only for clearer illustration) scale in D. Interestingly, these distributions are similar to coherence in Fig 3, reflecting that the coherence of alpha and theta bands are associated with phase locking rather than amplitude.

#### Theta PLV

The PLV analysis for stimulus-locked epochs again showed a right-lateralized increase in theta synchrony (Fig. 4C). And LI analysis indicated that this effect was significant in the time window from 375 ms to 420 ms *(p* < 0.05), for Aha as compared to Ctrl. For response-locked epochs, there was also a trend for a right-lateralized increase but the LI indices did not reach the significance level (Fig. 4D).

### Changes in alpha and beta power

#### Alpha power

In stimulus-locked epochs, there was no significant difference in alpha band (8–12 Hz) power between Aha and Ctrl conditions. In response-locked epochs, alpha power was found to be lower for Aha than for Ctrl trials. As shown in Figure 5, two clusters (obtained from cluster-based nonparametric randomization tests) of reduced alpha power were found (*p* < 0.05). One was in the frontal region, mostly in the left hemisphere, from 150 ms before response to response (−150 – 0 ms). In this cluster, alpha power was significantly decreased before the response. The second cluster was in the occipital region, from 335 ms before response to response (−335 – 0 ms). Interestingly, this cluster exhibited a gradual shift from the left to the right hemisphere with elapsing time. Before −220 ms, this cluster was mainly located in the left occipital area; after −220 ms, it moved to the right occipital and right superior parietal areas.

**Figure 5.**
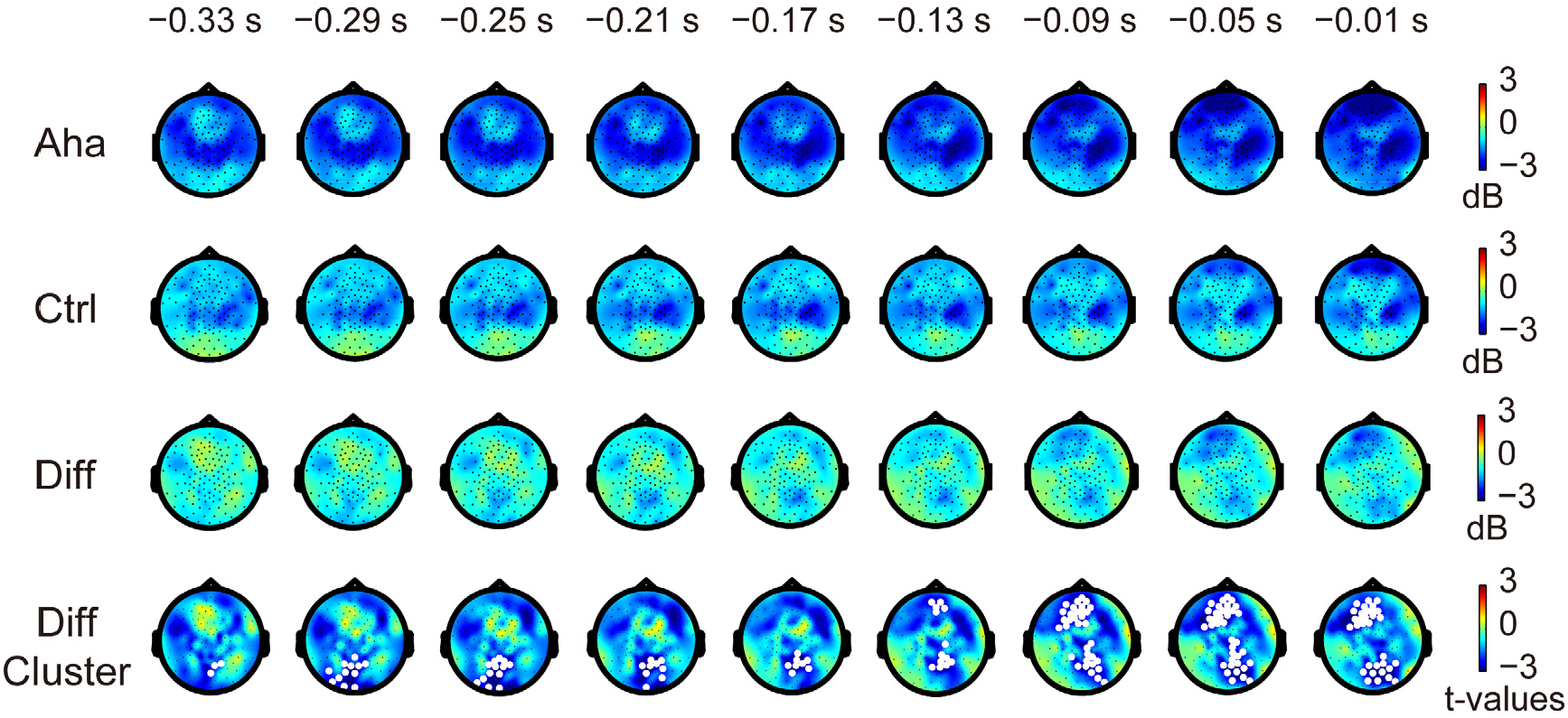
Changes in alpha power before the response to the Eureka inducing stimulus. The top row and second row: Amplitude (color scale) and distribution of alpha power (baseline corrected) in the Aha and Ctrl conditions at times (indicated above the snapshots) preceding the response. Third row: Difference between Aha and Ctrl (Aha - Ctrl). Fourth row: *t*-values of cluster-based nonparametric randomization test for the difference between Aha and Ctrl. The white regions represent clusters where alpha power decreased significantly *(p* < 0.05).

#### Beta Power

In stimulus-locked epochs, the Eureka effect was associated with a decrease in beta power (13–30 Hz) relative to control in the interval from 240 ms to 360 ms after stimulus onset *(p* < 0.05), as shown in Figure 6. This decrease was most prominent in the regions of the right parietal cortex and central gyrus, close to the midline. In response-locked epochs, the cluster-based nonparametric randomization test revealed a strong but not significant trend *(p* = 0.0789) of beta power decrease in the right parietal and right occipital cortices, in the time window of −380 – −285 ms before the response (Fig. 7).

**Figure 6.**
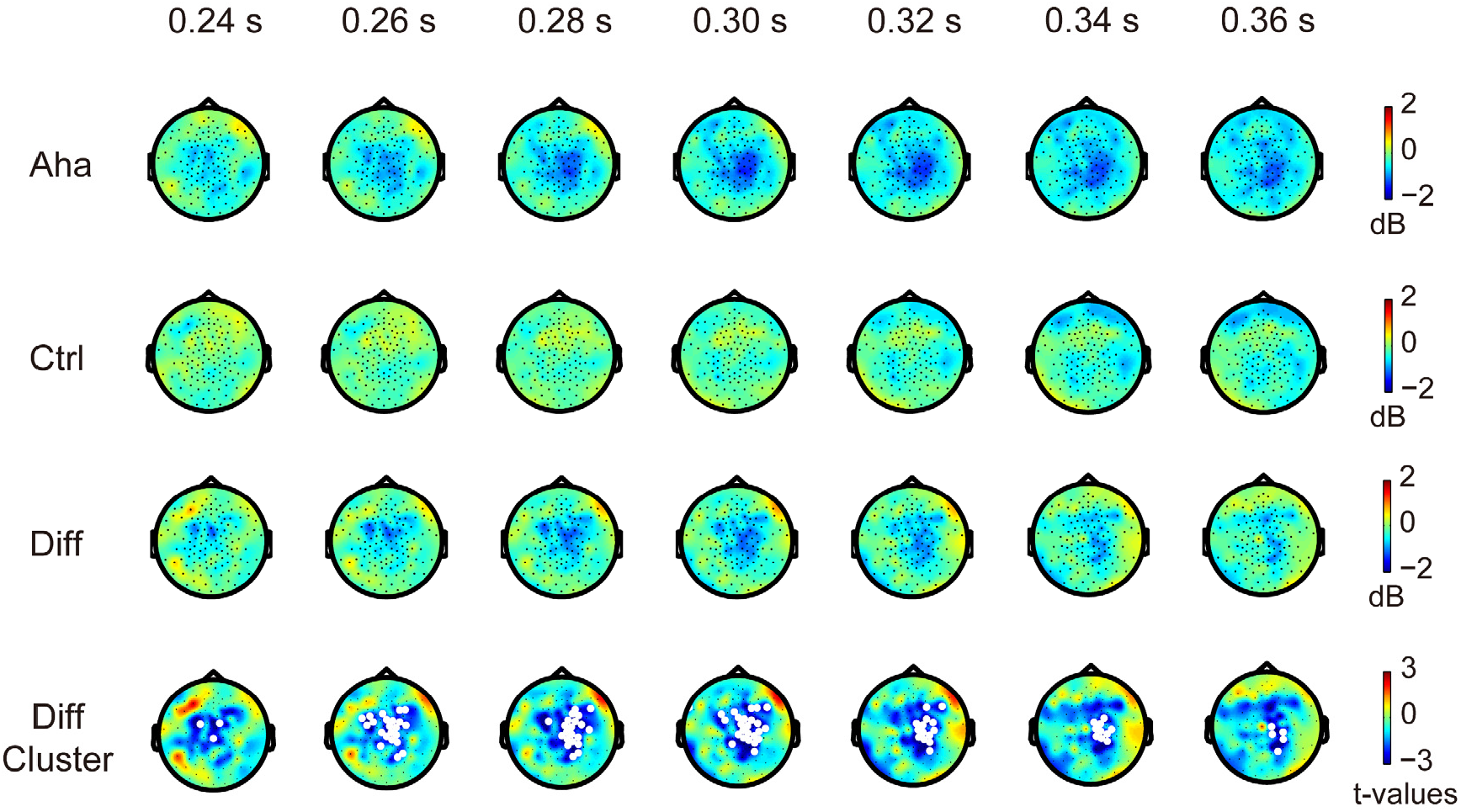
Changes in beta power after stimulus onset. Conventions as in Fig 5. Note the significant decrease in beta power (white regions in the bottom row) following the Eureka inducing stimulus.

**Figure 7.**
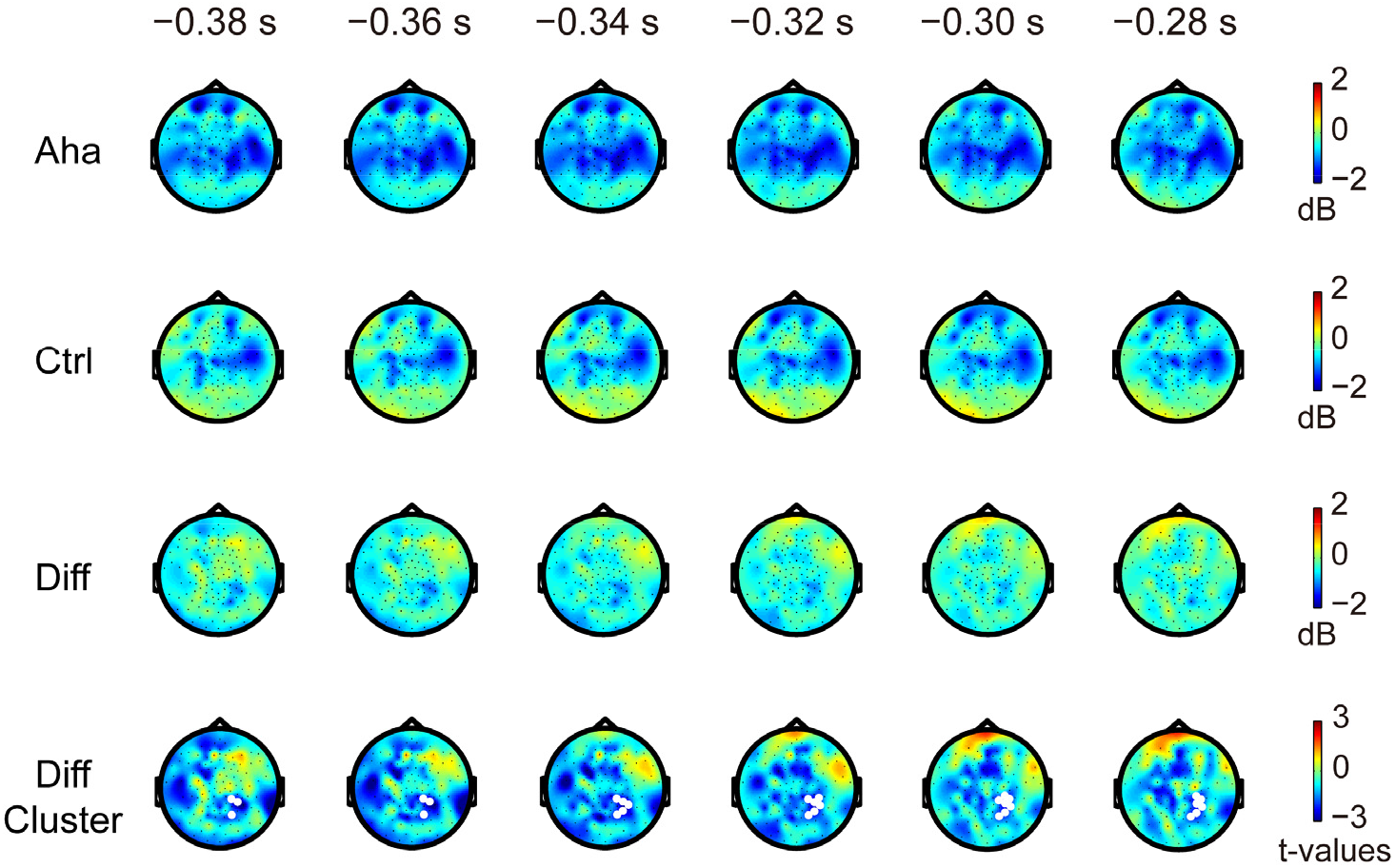
Changes in beta power before the response to the Eureka inducing stimulus. Conventions as in Figs. 5 and 6. Note the decrease in beta power (white regions in the bottom row; here *p* = 0.0789) found by cluster-based nonparametric randomization test for the difference between Aha and Ctrl, preceding the response to the Eureka inducing stimulus.

#### Theta power and Gamma Power

The power analysis of these frequency bands showed no significant differences between Eureka and controls.

### Changes in fractal dimensionality

We analyzed dimensionality for the time window from 0 to 500 ms after stimulus onset for 10 pre-defined clusters of electrodes (described in Materials and Methods). We observed significant reductions of dimensionality over right central and left parietal areas, and a significant increase over the right occipital area (Fig. 8A) in the Eureka trials. Furthermore, LI analysis showed that the dimensionality reduction was more pronounced (ln(*δ*) from 1.47 to 1.65, *p* < 0.05) in the temporal region of the right hemisphere (Fig. 8B). Similar results were obtained for response aligned activity patterns. In the time window from −500 ms to the response, there was a significant reduction of dimensionality over the right temporal area, and an increase of dimensionality over the left frontal area (Fig. 9A). The LI analysis confirmed the right lateralization of the dimensionality reduction (ln(*δ*) from 0.94 to 1.26, *p* < 0.05) over temporal areas (Fig. 9B).

**Figure 8.**
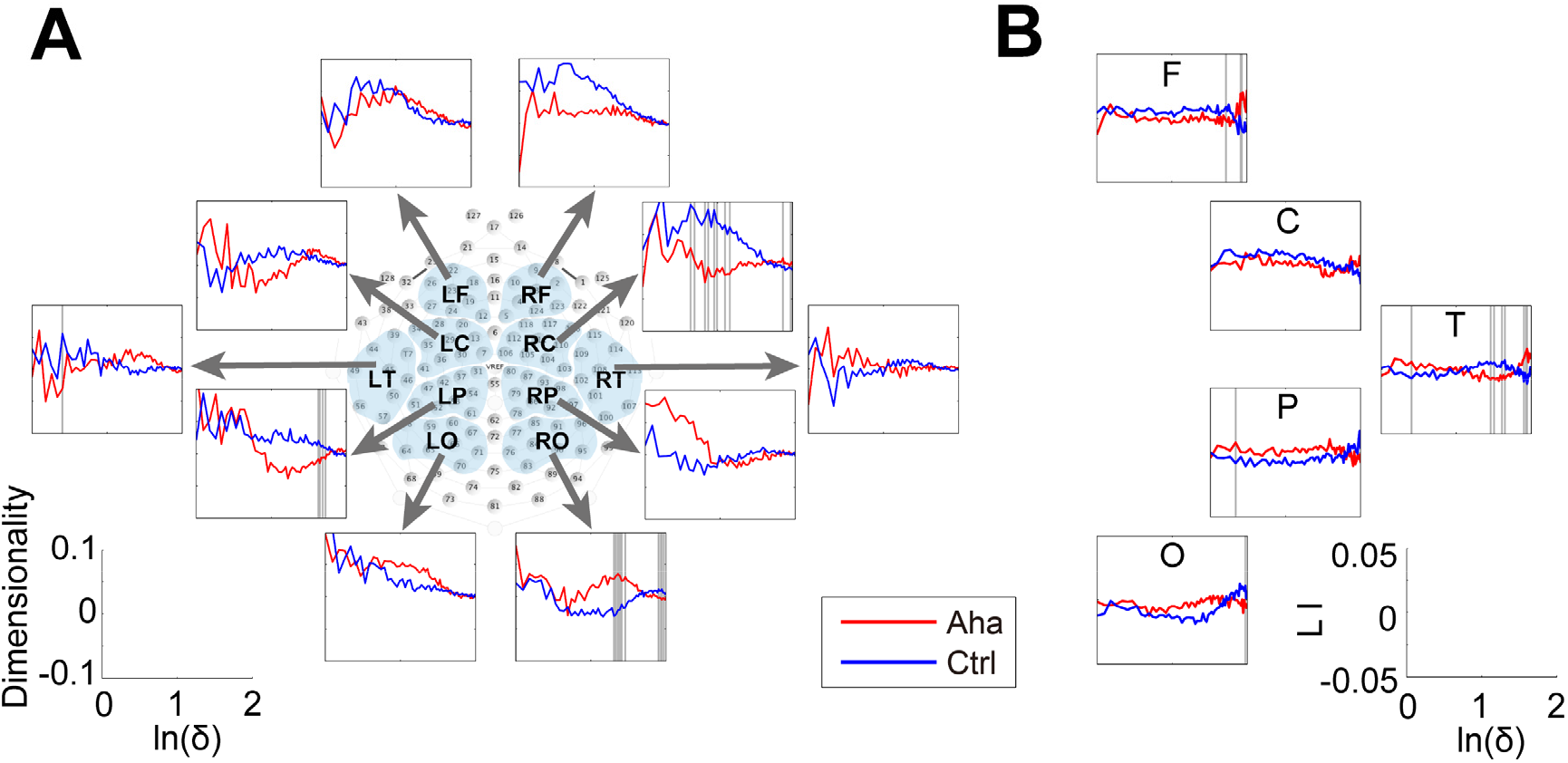
Changes in the dimensionality of activity vectors in the 500 ms interval following stimulus onset. (A) Panels representing the dimensionality of activity vectors derived from 10 clusters of recording sites as indicated by the map of electrode coverage (L=left, R=right; F=frontal, C=central, T=temporal, P=parietal, O=occipital). Baseline corrected. Ordinate: dimensionality; Abscissa: natural logarithm of scale size. (B) Laterality index (LI) of dimensionality for the clusters. Ordinate: laterality index; Abscissa: natural logarithm of scale size. The bins with significant differences between Aha and Ctrl are marked by grey lines.

**Figure 9.**
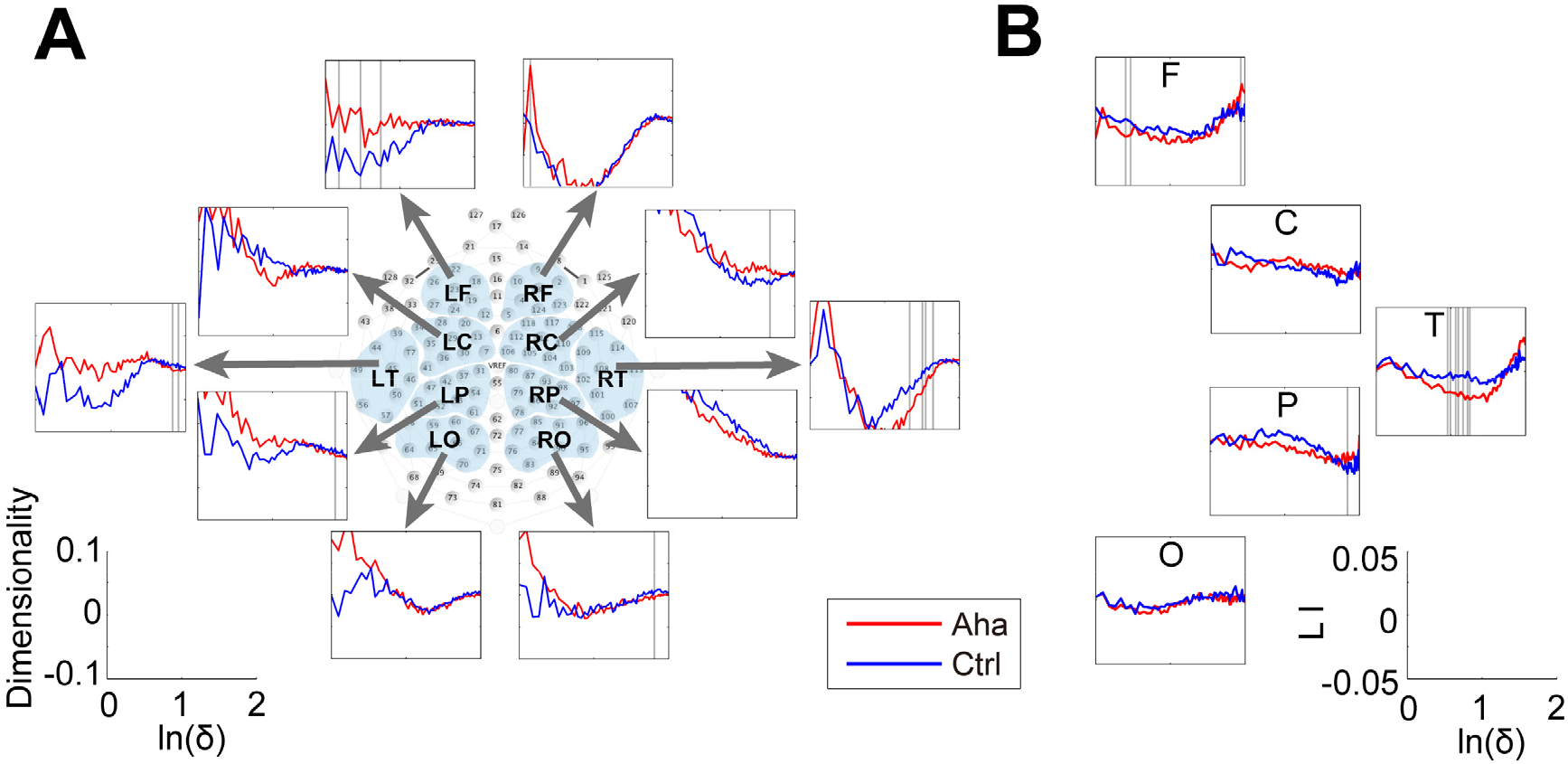
Changes in the dimensionality of activity vectors in the 500 ms interval before the response to the Eureka inducing stimulus. Conventions as in Fig. 8.

Taken together, in both the stimulus and response locked epochs the Eureka effect was associated with a reduction of fractal dimensionality in the right temporal cortex.

## Discussion

We investigated with EEG recordings brain activity during the Eureka effect and found that oscillatory activity in the alpha and theta bands exhibited right-hemisphere lateralized increases in coherence and phase locking, decreased power in the alpha band over frontal and occipital regions, and decreased power in the beta band over right parietal regions and the central gyrus, close to the midline. In addition, we observed a reduction of dimensionality for activity recorded from the frontal and temporal regions in the right hemisphere.

### Methodological considerations

A crucial question is whether our paradigm reliably elicited a Eureka effect. As reviewed in the introduction, the identification of neuronal correlates of the Eureka effect is hampered by the inability to precisely predict the time of its occurrence and to obtain sufficient trials per subject. Therefore, most of the classical conditions inducing Eureka effects are not suitable for neurobiological investigations. We have settled for a psychophysical task that allowed us to predict with some confidence, whether and when a stimulus would elicit a Eureka experience and to confirm its occurrence with a behavioral response. Subjects had to accomplish a difficult pattern completion task with physically identical stimuli that required flexible binding of features into a coherent gestalt and could be solved only in a minority of trials despite long inspection time when no additional cues were provided. However, by supplying additional associated information, subjects were enabled to solve the task promptly in the majority of trials, reporting sudden insight into the solution of the pattern completion problem (Dolan et al., 1997; Giovannelli et al., 2010). Debriefing after the experiment confirmed that subjects experienced a Eureka feeling at the *third stage* of the trials, in which they suddenly succeeded to identify the figure. We are thus confident, that the paradigm allowed for comparison of neuronal responses to physically identical stimuli that did or did not induce the Eureka effect. In addition, our design allowed us to run a large number of trials in the same subject, which is a prerequisite for neurophysiological studies.

Although our paradigm allowed narrowing the time interval during which the Eureka effect was bound to occur, response latencies were still variable. This could have been due to variable latencies of the Eureka effect or to variable lag times between the subjective Eureka experience and the motor response. In order to capture electrophysiological signatures of the Eureka state, we aligned for analysis neuronal responses to both stimulus and response onset, respectively. By adjusting the duration of the two analysis windows we made sure to cover the whole interval during which the Eureka effect was bound to occur. Because the results obtained for the two analysis windows were rather similar and located the effect to overlapping epochs between stimulus and response onset, we are confident that the electrophysiological changes reflect processes associated with the Eureka experience.

### Right-hemisphere lateralized alpha and theta activity

Our data indicate, that the Eureka effect is associated with distinct lateralized changes of coherence and power of oscillations in distinct frequency bands.

The most significant correlates of the Eureka effect were enhanced coherence and phase locking of alpha and theta oscillations over the right hemisphere. This enhanced synchronization was not associated with amplitude changes but only apparent in measures applied to sensor pairs. Hence our results indicate that for alpha and theta activities during the Eureka effect, the coherence is closely correlated with phase locking rather than amplitude.

Alpha oscillations have been associated with many different functions. The fact that they tend to be suppressed when neuronal circuits engage in information processing and high-frequency oscillations has been taken as indication that they represent an idling rhythm (Pfurtscheller et al., 1996; Jensen et al., 2002). However, there are also abundant indications for an involvement of alpha oscillations in active processing. Alpha oscillations have been suggested to serve the suppression of irrelevant information in tasks requiring focusing attention (Clarke et al., 2008; Zanto et al., 2010; Zanto et al., 2011; Klimesch, 2012), to coordinate widely distributed processes by playing the role of a carrier frequency for the establishment of coherence across various frequency bands (cross-frequency coupling) (Palva et al., 2005; Canolty and Knight, 2010; Jensen et al., 2014), to be involved in the maintenance of contents in working memory (Palva et al., 2005; Freunberger et al., 2009; Zanto et al., 2014; Foster et al., 2017; Riddle et al., 2020) and to mediate top-down control (Klimesch et al., 2010; Benedek et al., 2011; Palva and Palva, 2011; Samaha et al., 2015; Clayton et al., 2018). As elaborated further down we suggest that the association of increased alpha coherence and phase locking with the Eureka effect might be related to the maintenance and readout of contents from working memory and the top-down mediation of this information to facilitate scene segmentation and perceptual binding.

Theta band activities have been proposed to be involved in various memory processes such as memory encoding, maintenance, retrieval (Sauseng et al., 2010; Fell and Axmacher, 2011; Watrous et al., 2013; Burke et al., 2014; Roux and Uhlhaas, 2014; Herweg et al., 2020). Furthermore, the theta phase synchronization has been associated with various memory functions (Fell and Axmacher, 2011; Herweg et al., 2020). For example, investigators have observed increased theta phase locking (Liebe et al., 2012) and increased theta coherence (Sarnthein et al., 1998) between prefrontal and posterior recording sites, in working memory retention. Recently, increased theta phase locking has been found connecting a large set of brain regions to support memory encoding and recall (Burke et al., 2013; Clouter et al., 2017; Solomon et al., 2017; Wang et al., 2018; Solomon et al., 2019). In the present study, the prior experience stored in memory must be retrieved and used for perceptual grouping and recognition. We thus suggest that the increased theta coherence and phase locking might modulate the memory process in the Eureka effect.

An additional and non-exclusive possibility is that the enhanced right-lateralized alpha and theta phase synchronization is related to the “creative” processes associated with the Eureka effect. Similar electrographic signatures have been described in previous studies on creativity (Razoumnikova, 2000; Jung-Beeman et al., 2004; Runco, 2004; Grabner et al., 2007; Freunberger et al., 2008; Sandkuhler and Bhattacharya, 2008; Kounios and Beeman, 2014).

### Dissociation between alpha power and alpha phase locking

Somewhat unexpectedly there was a dissociation between alpha power and alpha phase locking. While alpha power decreased, coherence and phase locking increased. This excludes that enhanced coherence was simply a consequence of increased power and suggests non-trivial relations between the power of oscillatory activity and pair-wise synchrony. Power increases in oscillatory population responses can have two reasons. First, an increase in the number of neurons participating in the rhythmic activity, and second, enhanced precision of synchronization as this enhances the effective summation of currents. In the present case, we observed enhanced precision of synchronization as reflected by enhanced coherence and improved phase locking. This suggests that the Eureka effect was associated with a reduction of neurons engaged in alpha oscillations but at the same time with enhanced synchronization of the neuron populations participating in a right-hemisphere network engaged in alpha oscillations. A similar dissociation has been described by Freunberger et al. (2008), in an object recognition task with distorted color pictures. Associated with the recognition of the objects alpha power (9–13 Hz) decreased in occipital and right centro-temporal areas while phase locking (10–12 Hz) increased in a right-hemispheric long-range anterior-to-posterior network. In a later study on working memory, Freunberger et al. (2009) found in posterior areas (parietal-occipital) reduced alpha power and at the same time enhanced phase locking. These observations add to the notion that alpha oscillations reflect heterogeneous processes. In line with the idling hypothesis (Pfurtscheller et al., 1996; Jensen et al., 2002), the decrease in alpha power could reflect the increased engagement of right hemispheric networks and the simultaneous increase in synchronization the emergence of a specific, widely distributed but sparse network. The latter could serve the mediation of the top-down information required for the memory based perceptual closure of the Mooney images presented after the greyscale images. The involvement of alpha oscillations as a carrier frequency for top-down processes has been proposed (Palva et al., 2005; Canolty and Knight, 2010; Jensen et al., 2014). The increased alpha phase locking in the Eureka condition could thus be the reflection of the transient formation of a large scale coherent state comprising numerous areas in the right hemisphere.

### Right-hemisphere lateralized reduction of dimensionality

The interpretation that the Eureka effect is associated with the transient formation of a large but sparse right-hemispheric network oscillating in the alpha and theta frequency range is in agreement with the dimensionality reduction of the activity vector over the right hemisphere in the Eureka condition.

Dimensionality is a measure of the complexity of dynamic states. If dimensionality is high, coding space and the degrees of freedom are large but so is ambiguity. A reduction of dimensionality can be interpreted as a reduction of the number of possible states and hence as a reduction of ambiguity (Nikolic et al., 2008; Singer and Lazar, 2016). In the present experiments changes in dimensionality were in general variable and reached significance only rarely. However, there was a robust reduction of dimensionality over frontal and temporal areas of the right hemisphere. This agrees with the finding that the “solution” state leading to the Eureka experience was associated with enhanced coherence in the alpha and theta frequency bands, again over the right hemisphere. We interpret these changes as an indication that large cortical networks, in particular in the right hemisphere, converged towards a state identified as a solution state due to enhanced coherence and reduced variability.

### Decrease of beta activity

A transient decrease of beta activity has been reported in association with the transition from the maintenance of posture to movement initiation (Schoffelen et al., 2008; Pogosyan et al., 2009; Allen and MacKinnon, 2010; Kilavik et al., 2013; Armstrong et al., 2018) but it is unlikely that the observed decrease was related to the motor response of our subjects because it should have been canceled by subtraction of the control condition. A more likely interpretation is that the beta decrease is related to memory retrieval and the switching of cognitive states. The present task required retrieval of information from memory (Sheth et al., 2009; Kounios and Beeman, 2014) and engaging memory has been found to be associated with decreased beta power in the parietal and parieto-occipital areas (Pesonen et al., 2007; Sheth et al., 2009). As described in the study of Sheth et al. (2009) beta power also decreased in parietal, parieto-occipital, and centro-temporal areas in subjects solving verbal puzzles. Beta power also decreases transiently with switches in cognitive states (Okazaki et al., 2008; Engel and Fries, 2010; Piantoni et al., 2010) and with the disambiguation of visual stimuli (Minami et al., 2014). As sudden switches in cognitive states are a hallmark of the Eureka effect, we propose that the beta decrease observed in conjunction with the Eureka effect is related to such switches. To which extent the decrease in beta power is correlated to changes in neuronal activation cannot be inferred from EEG recordings. Two studies assessing neuronal activity with positron emission tomography (PET) and functional magnetic resonance imaging (fMRI) (Dolan et al., 1997; Eger et al., 2007) described an increase in activation in the parietal cortex associated with the Eureka experience. How this evidence relates to the transient decrease in beta power is unclear. As changes in neuronal dynamics such as increases and decreases of synchrony in particular frequency bands can occur without major changes in average firing rates, there need not be a conflict of these findings with our present results, both observations actually suggesting an involvement of the parietal cortex in the Eureka effect.

### The Eureka effect and gamma band

Although the processing of Mooney images, or in a more general context, perceptual closure has been shown to be associated with enhanced power and phase synchronization of gamma oscillations (Uhlhaas et al., 2006; Uhlhaas et al., 2009; Castelhano et al., 2013; Moratti et al., 2014), our data did not reveal any significant changes in the gamma band in the Eureka condition. This was unexpected as gamma synchronization is associated with binding operations (Singer, 1999) and successful matching of sensory evidence with contextual predictions in early visual areas (Peter et al., 2019). The solution of the detection task in our experiments did also require integration of sensory evidence with stored information. However, it is likely that these matching processes occurred at higher levels of the visual processing stream where oscillation frequencies are lower than in V1. Moreover, in EEG recordings the high-frequency oscillations in the gamma frequency range are detectable only when large assemblies of neurons get entrained in sustained, synchronized oscillations. Such entrainment can be achieved with high contrast drifting grating stimuli in low-level visual areas but not with cluttered scene stimuli as used in the present experiments (for review see Singer (2021)).

### Conclusions

Our data indicate that the Eureka effect involves primarily the right hemisphere in our sample of right-handed subjects. We interpret the increased coherence in the alpha and theta bands as indicators of the formation of a widely distributed network of cortical areas involved in the comparison of sensory evidence with information stored in working memory. The finding that enhanced coherence was associated with a reduction of dimensionality of the dynamic state but not with an increase in power in the respective frequency bands suggests as neuronal correlate of the Eureka experience the convergence of a large network to a dynamic state which is characterized by reduced dimensionality, reduced degrees of freedom and hence reduced ambiguity.

## Data and Code Availability Statement

The data used in this study are available via a request to the corresponding author upon signing a formal data sharing agreement. Codes used for this study are open to access at https://github.com/y-q-l/Mooney_EEG.

## Author contributions (CRediT)

**Yiqing Lu:** Conceptualization, Methodology, Investigation, Formal analysis, Data Curation, Visualization, Validation, Funding acquisition, Writing – Original draft, Writing – Review & Editing. **Wolf Singer:** Conceptualization, Methodology, Resources, Supervision, Funding acquisition, Writing – Review & Editing.

## Acknowledgements

This work was supported by scholarships from the Frankfurt Institute for Advanced Studies and the Chinese Scholarship Council (to Y.L.), by the Ernst Strüngmann Institute for Neuroscience in Cooperation with Max Planck Society, and by the Max Planck Institute for Brain Research. We would like to thank Gareth Bland for help on the script of dimensionality analysis.

## References

Ahissar M, Hochstein S (1997) Task difficulty and the specificity of perceptual learning. Nature 387:401–406. doi:10.1038/387401a0

Allen DP, MacKinnon CD (2010) Time-frequency analysis of movement-related spectral power in EEG during repetitive movements: a comparison of methods. J Neurosci Methods 186:107–115. doi:10.1016/j.jneumeth.2009.10.022

Amari S-i, Cichocki A, Yang HH (1996) A new learning algorithm for blind signal separation. Advances in neural information processing systems:757–763: Morgan Kaufmann Publishers. doi:10.5555/2998828.2998935

Anemuller J, Sejnowski TJ, Makeig S (2003) Complex independent component analysis of frequency-domain electroencephalographic data. Neural Netw 16:1311–1323. doi:10.1016/j.neunet.2003.08.003

Armstrong S, Sale MV, Cunnington R (2018) Neural Oscillations and the Initiation of Voluntary Movement. Front Psychol 9:2509. doi:10.3389/fpsyg.2018.02509

Avena-Koenigsberger A, Misic B, Sporns O (2017) Communication dynamics in complex brain networks. Nat Rev Neurosci 19:17–33. doi:10.1038/nrn.2017.149

Bell AJ, Sejnowski TJ (1995) An information-maximization approach to blind separation and blind deconvolution. Neural Comput 7:1129–1159. doi:10.1162/neco.1995.7.6.1129

Benedek M, Bergner S, Konen T, Fink A, Neubauer AC (2011) EEG alpha synchronization is related to top-down processing in convergent and divergent thinking. Neuropsychologia 49:3505–3511. doi:10.1016/j.neuropsychologia.2011.09.004

Bullmore E, Sporns O (2012) The economy of brain network organization. Nat Rev Neurosci 13:336–349. doi:10.1038/nrn3214

Burke JF, Zaghloul KA, Jacobs J, Williams RB, Sperling MR, Sharan AD, Kahana MJ (2013) Synchronous and asynchronous theta and gamma activity during episodic memory formation. J Neurosci 33:292–304. doi:10.1523/JNEUROSCI.2057-12.2013

Burke JF, Sharan AD, Sperling MR, Ramayya AG, Evans JJ, Healey MK, Beck EN, Davis KA, Lucas TH, 2nd, Kahana MJ (2014) Theta and high-frequency activity mark spontaneous recall of episodic memories. J Neurosci 34:11355–11365. doi:10.1523/JNEUROSCI.2654-13.2014

Canolty RT, Knight RT (2010) The functional role of cross-frequency coupling. Trends Cogn Sci 14:506–515. doi:10.1016/j.tics.2010.09.001

Castelhano J, Rebola J, Leitao B, Rodriguez E, Castelo-Branco M (2013) To Perceive or Not Perceive: The Role of Gamma-band Activity in Signaling Object Percepts. PLoS ONE 8. doi:10.1371/journal.pone.0066363

Clark A (2013) Whatever next? Predictive brains, situated agents, and the future of cognitive science. Behav Brain Sci 36:181–204. doi:10.1017/S0140525X12000477

Clarke AR, Barry RJ, Heaven PC, McCarthy R, Selikowitz M, Byrne MK (2008) EEG coherence in adults with attention-deficit/hyperactivity disorder. Int J Psychophysiol 67:35–40. doi:10.1016/j.ijpsycho.2007.10.001

Clayton MS, Yeung N, Cohen Kadosh R (2018) The many characters of visual alpha oscillations. Eur J Neurosci 48:2498–2508. doi:10.1111/ejn.13747

Clouter A, Shapiro KL, Hanslmayr S (2017) Theta Phase Synchronization Is the Glue that Binds Human Associative Memory. Curr Biol 27:3143–3148 e3146. doi:10.1016/j.cub.2017.09.001

de Lange FP, Heilbron M, Kok P (2018) How Do Expectations Shape Perception? Trends Cogn Sci 22:764–779. doi:10.1016/j.tics.2018.06.002

Dolan RJ, Fink GR, Rolls E, Booth M, Holmes A, Frackowiak RS, Friston KJ (1997) How the brain learns to see objects and faces in an impoverished context. Nature 389:596–599. doi:10.1038/39309

Eger E, Henson RN, Driver J, Dolan RJ (2007) Mechanisms of top-down facilitation in perception of visual objects studied by FMRI. Cereb Cortex 17:2123–2133. doi:10.1093/cercor/bhl119

Engel AK, Fries P (2010) Beta-band oscillations--signalling the status quo? Curr Opin Neurobiol 20:156–165. doi:10.1016/j.conb.2010.02.015

Fell J, Axmacher N (2011) The role of phase synchronization in memory processes. Nat Rev Neurosci 12:105–118. doi:10.1038/nrn2979

Foster JJ, Bsales EM, Jaffe RJ, Awh E (2017) Alpha-Band Activity Reveals Spontaneous Representations of Spatial Position in Visual Working Memory. Curr Biol 27:3216–3223 e3216. doi:10.1016/j.cub.2017.09.031

Freunberger R, Klimesch W, Griesmayr B, Sauseng P, Gruber W (2008) Alpha phase coupling reflects object recognition. Neuroimage 42:928–935. doi:10.1016/j.neuroimage.2008.05.020

Freunberger R, Fellinger R, Sauseng P, Gruber W, Klimesch W (2009) Dissociation between phase-locked and nonphase-locked alpha oscillations in a working memory task. Hum Brain Mapp 30:3417–3425. doi:10.1002/hbm.20766

Fries P (2009) Neuronal gamma-band synchronization as a fundamental process in cortical computation. Annu Rev Neurosci 32:209–224. doi:10.1146/annurev.neuro.051508.135603

Friston KJ, Stephan KE (2007) Free-energy and the brain. Synthese 159:417–458. doi:10.1007/s11229-007-9237-y

Giovannelli F, Silingardi D, Borgheresi A, Feurra M, Amati G, Pizzorusso T, Viggiano MP, Zaccara G, Berardi N, Cincotta M (2010) Involvement of the parietal cortex in perceptual learning (Eureka effect): an interference approach using rTMS. Neuropsychologia 48:1807–1812. doi:10.1016/j.neuropsychologia.2010.02.031

Goold JE, Meng M (2016) Visual Search of Mooney Faces. Front Psychol 7:155. doi:10.3389/fpsyg.2016.00155

Gorlin S, Meng M, Sharma J, Sugihara H, Sur M, Sinha P (2012) Imaging prior information in the brain. Proc Natl Acad Sci U S A 109:7935–7940. doi:10.1073/pnas.1111224109

Grabner RH, Fink A, Neubauer AC (2007) Brain correlates of self-rated originality of ideas: evidence from event-related power and phase-locking changes in the EEG. Behav Neurosci 121:224–230. doi:10.1037/0735-7044.121.1.224

Grassberger P, Procaccia I (2004) Measuring the strangeness of strange attractors. In: The Theory of Chaotic Attractors, pp 170–189: Springer. doi:10.1007/978-0-387-21830-4_12

Gray CM, Konig P, Engel AK, Singer W (1989) Oscillatory Responses in Cat Visual-Cortex Exhibit Inter-Columnar Synchronization Which Reflects Global Stimulus Properties. Nature 338:334–337. doi:10.1038/338334a0

Griffin G, Holub A, Perona P (2007) Caltech-256 object category dataset.

Grutzner C, Uhlhaas PJ, Genc E, Kohler A, Singer W, Wibral M (2010) Neuroelectromagnetic correlates of perceptual closure processes. J Neurosci 30:8342–8352. doi:10.1523/JNEUROSCI.5434-09.2010

Haddad RA, Akansu AN (1991) A class of fast Gaussian binomial filters for speech and image processing. IEEE Transactions on Signal Processing 39:723–727. doi:10.1109/78.80892

Herweg NA, Solomon EA, Kahana MJ (2020) Theta Oscillations in Human Memory. Trends Cogn Sci 24:208–227. doi:10.1016/j.tics.2019.12.006

Hipp JF, Engel AK, Siegel M (2011) Oscillatory synchronization in large-scale cortical networks predicts perception. Neuron 69:387–396. doi:10.1016/j.neuron.2010.12.027

Hsieh PJ, Vul E, Kanwisher N (2010) Recognition alters the spatial pattern of FMRI activation in early retinotopic cortex. J Neurophysiol 103:1501–1507. doi:10.1152/jn.00812.2009

Jansen A, Menke R, Sommer J, Forster AF, Bruchmann S, Hempleman J, Weber B, Knecht S (2006) The assessment of hemispheric lateralization in functional MRI--robustness and reproducibility. Neuroimage 33:204–217. doi:10.1016/j.neuroimage.2006.06.019

Jensen O, Gelfand J, Kounios J, Lisman JE (2002) Oscillations in the alpha band (9-12 Hz) increase with memory load during retention in a short-term memory task. Cereb Cortex 12:877–882. doi:10.1093/cercor/12.8.877

Jensen O, Gips B, Bergmann TO, Bonnefond M (2014) Temporal coding organized by coupled alpha and gamma oscillations prioritize visual processing. Trends Neurosci 37:357–369. doi:10.1016/j.tins.2014.04.001

Jung-Beeman M, Bowden EM, Haberman J, Frymiare JL, Arambel-Liu S, Greenblatt R, Reber PJ, Kounios J (2004) Neural activity when people solve verbal problems with insight. PLoS Biol 2:E97. doi:10.1371/journal.pbio.0020097

Kaplan CA, Simon HA (1990) In search of insight. Cognitive psychology 22:374–419. doi:10.1016/0010-0285(90)90008-R

Kersten D, Mamassian P, Yuille A (2004) Object perception as Bayesian inference. Annu Rev Psychol 55:271–304. doi:10.1146/annurev.psych.55.090902.142005

Kilavik BE, Zaepffel M, Brovelli A, MacKay WA, Riehle A (2013) The ups and downs of beta oscillations in sensorimotor cortex. Exp Neurol 245:15–26. doi:10.1016/j.expneurol.2012.09.014

Klimesch W (2012) alpha-band oscillations, attention, and controlled access to stored information. Trends Cogn Sci 16:606–617. doi:10.1016/j.tics.2012.10.007

Klimesch W, Freunberger R, Sauseng P (2010) Oscillatory mechanisms of process binding in memory. Neurosci Biobehav Rev 34:1002–1014. doi:10.1016/j.neubiorev.2009.10.004

Kounios J, Beeman M (2014) The cognitive neuroscience of insight. Annu Rev Psychol 65:71–93. doi:10.1146/annurev-psych-010213-115154

Lachaux JP, Rodriguez E, Martinerie J, Varela FJ (1999) Measuring phase synchrony in brain signals. Hum Brain Mapp 8:194–208. doi:10.1002/(sici)1097-0193(1999)8:4<194::aid-hbm4>3.0.co;2-c

Lee T-W, Girolami M, Sejnowski TJ (1999) Independent component analysis using an extended infomax algorithm for mixed subgaussian and supergaussian sources. Neural Comput 11:417–441. doi:10.1162/089976699300016719

Liebe S, Hoerzer GM, Logothetis NK, Rainer G (2012) Theta coupling between V4 and prefrontal cortex predicts visual short-term memory performance. Nat Neurosci 15:456–462, S451-452. doi:10.1038/nn.3038

Lim JS (1990) Two-dimensional signal and image processing. Englewood Cliffs, NJ, Prentice Hall.

Marie C, Trainor LJ (2013) Development of simultaneous pitch encoding: infants show a high voice superiority effect. Cereb Cortex 23:660–669. doi:10.1093/cercor/bhs050

Maris E, Oostenveld R (2007) Nonparametric statistical testing of EEG- and MEG-data. J Neurosci Methods 164:177–190. doi:10.1016/j.jneumeth.2007.03.024

McKeeff TJ, Tong F (2007) The timing of perceptual decisions for ambiguous face stimuli in the human ventral visual cortex. Cereb Cortex 17:669–678. doi:10.1093/cercor/bhk015

Minami T, Noritake Y, Nakauchi S (2014) Decreased beta-band activity is correlated with disambiguation of hidden figures. Neuropsychologia 56:9–16. doi:10.1016/j.neuropsychologia.2013.12.026

Moratti S, Mendez-Bertolo C, Del-Pozo F, Strange BA (2014) Dynamic gamma frequency feedback coupling between higher and lower order visual cortices underlies perceptual completion in humans. Neuroimage 86:470–479. doi:10.1016/j.neuroimage.2013.10.037

Nikolic D, Moca VV, Singer W, Muresan RC (2008) Properties of multivariate data investigated by fractal dimensionality. Journal of Neuroscience Methods 172:27–33. doi:10.1016/j.jneumeth.2008.04.007

Okazaki M, Kaneko Y, Yumoto M, Arima K (2008) Perceptual change in response to a bistable picture increases neuromagnetic beta-band activities. Neurosci Res 61:319–328. doi:10.1016/j.neures.2008.03.010

Oostenveld R, Fries P, Maris E, Schoffelen JM (2011) FieldTrip: Open source software for advanced analysis of MEG, EEG, and invasive electrophysiological data. Comput Intell Neurosci 2011:156869. doi:10.1155/2011/156869

Palva JM, Palva S, Kaila K (2005) Phase synchrony among neuronal oscillations in the human cortex. J Neurosci 25:3962–3972. doi:10.1523/JNEUROSCI.4250-04.2005

Palva S, Palva JM (2011) Functional roles of alpha-band phase synchronization in local and large-scale cortical networks. Front Psychol 2:204. doi:10.3389/fpsyg.2011.00204

Pesonen M, Hamalainen H, Krause CM (2007) Brain oscillatory 4-30 Hz responses during a visual n-back memory task with varying memory load. Brain Res 1138:171–177. doi:10.1016/j.brainres.2006.12.076

Peter A, Uran C, Klon-Lipok J, Roese R, van Stijn S, Barnes W, Dowdall JR, Singer W, Fries P, Vinck M (2019) Surface color and predictability determine contextual modulation of V1 firing and gamma oscillations. eLife 8:e42101. doi:10.7554/eLife.42101

Pfurtscheller G, Stancak A, Jr., Neuper C (1996) Event-related synchronization (ERS) in the alpha band--an electrophysiological correlate of cortical idling: a review. Int J Psychophysiol 24:39–46. doi:10.1016/s0167-8760(96)00066-9

Piantoni G, Kline KA, Eagleman DM (2010) Beta oscillations correlate with the probability of perceiving rivalrous visual stimuli. J Vis 10:18. doi:10.1167/10.13.18

Pogosyan A, Gaynor LD, Eusebio A, Brown P (2009) Boosting cortical activity at Beta-band frequencies slows movement in humans. Curr Biol 19:1637–1641. doi:10.1016/j.cub.2009.07.074

Razoumnikova OM (2000) Functional organization of different brain areas during convergent and divergent thinking: an EEG investigation. Brain Res Cogn Brain Res 10:11–18. doi:10.1016/s0926-6410(00)00017-3

Riddle J, Scimeca JM, Cellier D, Dhanani S, D’Esposito M (2020) Causal Evidence for a Role of Theta and Alpha Oscillations in the Control of Working Memory. Curr Biol 30:1748–1754 e1744. doi:10.1016/j.cub.2020.02.065

Rosenberg JR, Amjad AM, Breeze P, Brillinger DR, Halliday DM (1989) The Fourier approach to the identification of functional coupling between neuronal spike trains. Prog Biophys Mol Biol 53:1–31. doi:10.1016/0079-6107(89)90004-7

Roux F, Uhlhaas PJ (2014) Working memory and neural oscillations: alpha-gamma versus theta-gamma codes for distinct WM information? Trends Cogn Sci 18:16–25. doi:10.1016/j.tics.2013.10.010

Runco MA (2004) Creativity. Annu Rev Psychol 55:657–687. doi:10.1146/annurev.psych.55.090902.141502

Samaha J, Bauer P, Cimaroli S, Postle BR (2015) Top-down control of the phase of alpha-band oscillations as a mechanism for temporal prediction. Proc Natl Acad Sci U S A 112:8439–8444. doi:10.1073/pnas.1503686112

Sandkuhler S, Bhattacharya J (2008) Deconstructing insight: EEG correlates of insightful problem solving. PLoS ONE 3:e1459. doi:10.1371/journal.pone.0001459

Sarnthein J, Petsche H, Rappelsberger P, Shaw GL, von Stein A (1998) Synchronization between prefrontal and posterior association cortex during human working memory. Proc Natl Acad Sci U S A 95:7092–7096. doi:10.1073/pnas.95.12.7092

Sauseng P, Griesmayr B, Freunberger R, Klimesch W (2010) Control mechanisms in working memory: a possible function of EEG theta oscillations. Neurosci Biobehav Rev 34:1015–1022. doi:10.1016/j.neubiorev.2009.12.006

Schoffelen JM, Oostenveld R, Fries P (2008) Imaging the human motor system’s beta-band synchronization during isometric contraction. Neuroimage 41:437–447. doi:10.1016/j.neuroimage.2008.01.045

Seghier ML (2008) Laterality index in functional MRI: methodological issues. Magn Reson Imaging 26:594–601. doi:10.1016/j.mri.2007.10.010

Sehatpour P, Molholm S, Schwartz TH, Mahoney JR, Mehta AD, Javitt DC, Stanton PK, Foxe JJ (2008) A human intracranial study of long-range oscillatory coherence across a frontal-occipital-hippocampal brain network during visual object processing. Proc Natl Acad Sci U S A 105:4399–4404. doi:10.1073/pnas.0708418105

Sheth BR, Sandkuhler S, Bhattacharya J (2009) Posterior Beta and anterior gamma oscillations predict cognitive insight. J Cogn Neurosci 21:1269–1279. doi:10.1162/jocn.2009.21069

Singer W (1999) Neuronal synchrony: a versatile code for the definition of relations? Neuron 24:49–65, 111-125. doi:10.1016/s0896-6273(00)80821-1

Singer W (2018) Neuronal oscillations: unavoidable and useful? Eur J Neurosci 48:2389–2398. doi:10.1111/ejn.13796

Singer W (2021) Recurrent dynamics in the cerebral cortex: integration of sensory evidence with stored knowledge. in review.

Singer W, Gray CM (1995) Visual feature integration and the temporal correlation hypothesis. Annu Rev Neurosci 18:555–586. doi:10.1146/annurev.ne.18.030195.003011

Singer W, Lazar A (2016) Does the Cerebral Cortex Exploit High-Dimensional, Non-linear Dynamics for Information Processing? Front Comput Neurosci 10:99. doi:10.3389/fncom.2016.00099

Solomon EA, Stein JM, Das S, Gorniak R, Sperling MR, Worrell G, Inman CS, Tan RJ, Jobst BC, Rizzuto DS, Kahana MJ (2019) Dynamic Theta Networks in the Human Medial Temporal Lobe Support Episodic Memory. Curr Biol 29:1100–1111 e1104. doi:10.1016/j.cub.2019.02.020

Solomon EA, Kragel JE, Sperling MR, Sharan A, Worrell G, Kucewicz M, Inman CS, Lega B, Davis KA, Stein JM, Jobst BC, Zaghloul KA, Sheth SA, Rizzuto DS, Kahana MJ (2017) Widespread theta synchrony and high-frequency desynchronization underlies enhanced cognition. Nat Commun 8:1704. doi:10.1038/s41467-017-01763-2

Sporns O, Chialvo DR, Kaiser M, Hilgetag CC (2004) Organization, development and function of complex brain networks. Trends Cogn Sci 8:418–425. doi:10.1016/j.tics.2004.07.008

Sprugnoli G, Rossi S, Emmerdorfer A, Rossi A, Liew S-L, Tatti E, di Lorenzo G, Pascual-Leone A, Santarnecchi E (2017) Neural correlates of Eureka moment. Intelligence. doi:10.1016/j.intell.2017.03.004

Srinath R, Ray S (2014) Effect of amplitude correlations on coherence in the local field potential. J Neurophysiol 112:741–751. doi:10.1152/jn.00851.2013

Sternberg RJ, Davidson JE (1995) The nature of insight: The MIT Press.

Tallon-Baudry C, Bertrand O (1999) Oscillatory gamma activity in humans and its role in object representation. Trends Cogn Sci 3:151–162. doi:10.1016/s1364-6613(99)01299-1

Tovee MJ, Rolls ET, Ramachandran VS (1996) Rapid visual learning in neurones of the primate temporal visual cortex. Neuroreport 7:2757–2760. doi:10.1097/00001756-199611040-00070

Uhlhaas PJ, Roux F, Singer W, Haenschel C, Sireteanu R, Rodriguez E (2009) The development of neural synchrony reflects late maturation and restructuring of functional networks in humans. Proc Natl Acad Sci U S A 106:9866–9871. doi:10.1073/pnas.0900390106

Uhlhaas PJ, Linden DE, Singer W, Haenschel C, Lindner M, Maurer K, Rodriguez E (2006) Dysfunctional long-range coordination of neural activity during Gestalt perception in schizophrenia. J Neurosci 26:8168–8175. doi:10.1523/JNEUROSCI.2002-06.2006

Volberg G, Wutz A, Greenlee MW (2013) Top-down control in contour grouping. PLoS One 8:e54085. doi:10.1371/journal.pone.0054085

Wang D, Clouter A, Chen Q, Shapiro KL, Hanslmayr S (2018) Single-Trial Phase Entrainment of Theta Oscillations in Sensory Regions Predicts Human Associative Memory Performance. J Neurosci 38:6299–6309. doi:10.1523/JNEUROSCI.0349-18.2018

Watrous AJ, Tandon N, Conner CR, Pieters T, Ekstrom AD (2013) Frequency-specific network connectivity increases underlie accurate spatiotemporal memory retrieval. Nat Neurosci 16:349–356. doi:10.1038/nn.3315

Wilke M, Schmithorst VJ (2006) A combined bootstrap/histogram analysis approach for computing a lateralization index from neuroimaging data. Neuroimage 33:522–530. doi:10.1016/j.neuroimage.2006.07.010

Zanto TP, Chadick JZ, Gazzaley A (2014) Anticipatory alpha phase influences visual working memory performance. Neuroimage 85 Pt 2:794–802. doi:10.1016/j.neuroimage.2013.07.048

Zanto TP, Rubens MT, Bollinger J, Gazzaley A (2010) Top-down modulation of visual feature processing: the role of the inferior frontal junction. Neuroimage 53:736–745. doi:10.1016/j.neuroimage.2010.06.012

Zanto TP, Rubens MT, Thangavel A, Gazzaley A (2011) Causal role of the prefrontal cortex in top-down modulation of visual processing and working memory. Nat Neurosci 14:656–661. doi:10.1038/nn.2773

