## Supplementary Material for "Dynamic signatures of the Eureka effect: An EEG study"

Yiqing Lu [a, b, c, d] * and Wolf Singer [a, b, c]

[a] Ernst Strüngmann Institute for Neuroscience in Cooperation with Max Planck Society, 60528 Frankfurt am Main, Germany

[b] Frankfurt Institute for Advanced Studies, 60438 Frankfurt am Main, Germany

[c] Department of Neurophysiology, Max Planck Institute for Brain Research, 60528 Frankfurt am Main, Germany

[d] Department of Biology, Technische Universität Darmstadt, 64287 Darmstadt, Germany

### The selection of the appropriate low-pass spatial frequency for the stimuli presented to induce the Eureka effect

**Visual stimuli.** The greyscale images were low pass filtered at different spatial frequencies to produce Mooney images with different degradation levels and then tested for the degree to which they were recognizable. For a series of Mooney images derived from one original image, the difficulty to recognize the images increased to the extent that low spatial frequencies were cut off. Recognition thresholds were determined by presenting sequences of degraded images, starting with the least recognizable. Each trial contained 8 Mooney images (derived from one original image) with cut-off frequencies from 10 Hz to 80 Hz (Fig. S1).

**Behavioral task.** Each Mooney image was presented for 2 seconds, at the center of the screen. If the subject could not identify the object within 2 s (the button was not pressed), the next image would appear (i.e., if the subject could not identify the 10 Hz object, then the 20 Hz object would appear; then 30 Hz, 40 Hz, until the last one at 80 Hz). Whenever the subject could identify the object in the image, she/he had to press the 'Yes' button as soon as possible. If the subject was unable to identify the 80 Hz object, the trial was aborted. Figure S2 shows the task procedure.

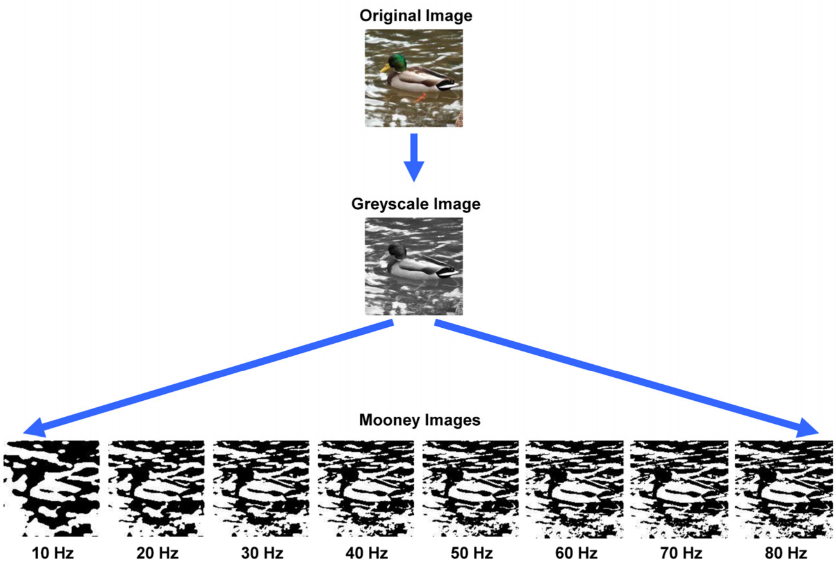


**Figure S1. The procedure of image manipulation.** The greyscale and two-tone Mooney images are derived from the top original image. The bottom row represents the images filtered with low-pass cut-off frequencies from 10 Hz (left) to 80 Hz (right).

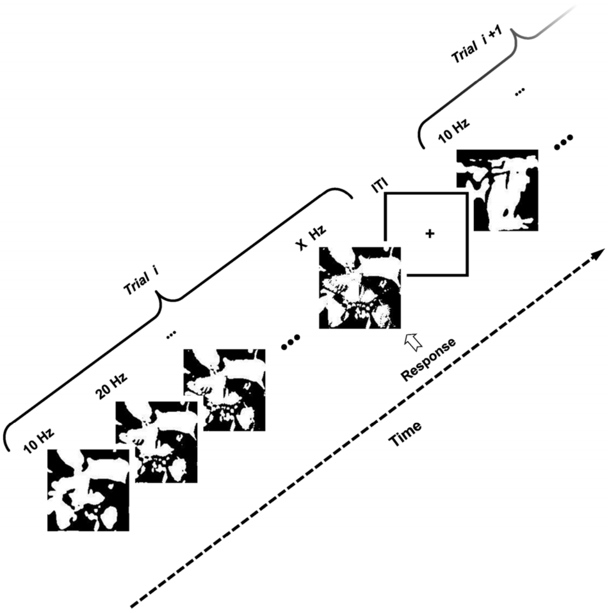


**Figure S2. The behavioral task for the selection of the cut-off frequency used in the main experiment.** One trial consisted of the presentation of a sequence of images with ascending cut-off frequency (from 10 to 80 Hz). Each image was shown for 2 sec, the intertrial interval (ITI) was 1 sec. Subjects were asked to respond swiftly once they had recognized the object (it is a butterfly in this example). The “X Hz” in the figure indicates the cut-off frequency at which the subject could recognize the object.

**Participants.** Ten healthy subjects (Age 25.3 ± 1.3 y, 5 males, 5 females) were recruited. In the behavioral analysis, all of the ten subjects were included. None of these subjects took part in the main experiment with EEG recordings. All subjects were naïve to the experiment, were right-handed, had normal or corrected-to-normal vision, and had no history of neurological or psychiatric disorders. They gave written informed consent before the experiment. The study was approved by the ethical committee of the Goethe University, Frankfurt, and was conducted in accordance with the Declaration of Helsinki. The subjects were recruited from local universities and got paid 15 Euros per hour for their participation.

**Behavioral results.** The recognition rates for the various degradation levels were 3.1%, 10.6%, 8.5%, 8.8%, 7.6%, 6.4%, 6.2%, and 4.1%, for cut-off frequencies ranging from 10 Hz to 80 Hz. The cumulative recognition rate (CRR) represents the theoretical recognition rate on a specific degradation level for naïve Mooney images of our image database. CRRs were calculated from 10 Hz to 80 Hz as well (Fig. S3). Additionally, in 44.7% of the trials subjects failed to respond, i.e., they could not recognize (NR) these images. The sum of CRR at 80 Hz and NR represents all trials, i.e., 100% (Fig. S3).

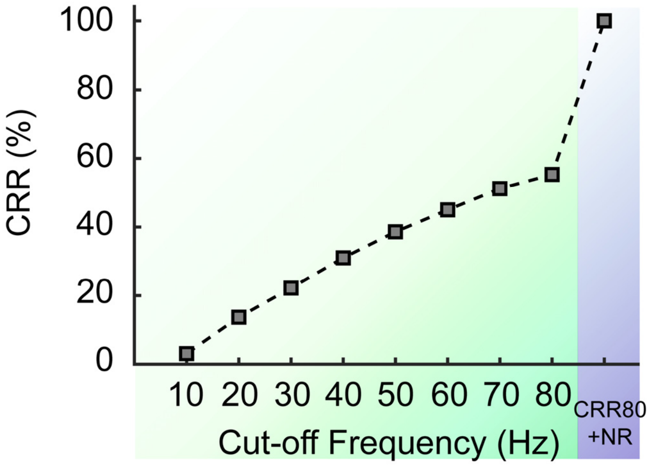


**Figure S3. The cumulative recognition rate (CRR)** to images with different degradation levels, the cut-off frequency from 10 Hz to 80 Hz (Green area). **The sum of CRR at 80 Hz and "not recognize" (NR)** (CRR80+NR, purple area).

**The selection of cut-off frequencies for the main experiment.** There are two conflicting requirements for the selection of the suitable cut-off frequencies. On the one hand, the identification must be difficult enough to require cognitive resources, on the other hand, the subjects must experience sufficient success to remain motivated and to experience the Eureka effect. According to the behavioral results, 3.1% of the images can be recognized with the 10 Hz cut-off frequency, which we considered to be too low to maintain high attention. With the 20 Hz cut-off frequency 10.6% of the images could be recognized (the former 3.1% of the images are not included). Because the images with 20 Hz cut-off frequency are easier to identify than those with 10 Hz cut-off frequency, we assumed that the former 3.1% of the images can also be recognized with 20 Hz cut-off frequency. So, in total 3.1% + 10.6% = 13.7%, i.e., the CRR at 20 Hz, of the images should have been identifiable with a 20 Hz cut-off frequency, implying that subjects can recognize one image out of approximately seven images (13.7%). This should have been sufficient to maintain the attention of subjects while being difficult enough to produce a Eureka effect. Therefore, we opted for a 20 Hz cut-off frequency for all stimuli presented in the main experiment.
